# Regulation of polyphosphate glucokinase gene expression through co-transcriptional processing in *Mycobacterium tuberculosis* H37Rv

**DOI:** 10.1101/2020.03.31.018051

**Authors:** Naveen Prakash Bokolia, Inshad Ali Khan

**Author notes:** Tel: []; Fax: [91-191- 2569333], ] (Inshad Ali Khan).

## Abstract

Transcription is the process that allows the simultaneous folding of RNA molecules, known as co-transcriptional folding. This folding determines the functional properties of RNA molecules and possibly having a critical role during the synthesis as well. This functioning includes the characterized properties of riboswitches and ribozymes as well, which is significant when the transcription rate is comparable to the cellular environment. This study aims to discover a novel non-coding region that is important in the genetic expression of *Mycobacterium tuberculosis*. In this work, we identified a novel non-coding element of polyphosphate glucokinase (*ppgk*) gene that undergoes cleavage activity during the transcriptional process in *Mycobacterium tuberculosis*. We revealed that cleavage occurs within the nascent RNA, and the resultant cleaved 3’RNA fragment carries the Shine-Dalgarno (SD) sequence and expression platform. Site-specific mutations provide a strong correlation between the disruption of cleavage activity and expression of *ppgk* gene. We concluded that co-transcriptional processing at the noncoding region as the required mechanism for *ppgk* expression that remains constitutive within the bacterial environment. The underlying reason for *ppgk* mRNA processing and expression is correlated because the non-coding counterpart adopts a hairpin domain that sequesters ribosomal binding site. Thus, the mRNA processing at the immediate upstream of Shine-Dalgarno sequence is required to prevent this sequestration and subsequent expression as well. This study defines the molecular mechanism that is dependent on the transient but highly active structural features of the nascent RNA.

## Background

*Mycobacterium tuberculosis* is the causative agent of tuberculosis in the human population (Smith 2003). Apart from slow-growing features, this organism has a high adaptation to survive under stress, specifically during infection in macrophages (Cook et al. 2009). There are several challenges in stepping for understanding the genetic regulation of *M. tuberculosis*. However, many studies have become successful in the identification of critical regulatory elements (Rodriguez et al. 2002; Manganelli et al. 2004; Raman et al. 2004; Turkarslan et al. 2015).

Within the last few years, newer perspectives of noncoding RNA elements have provided insightful directions in understanding gene regulation. Riboswitches are the most recognized elements that involve direct binding of the respective metabolite and, in consequence, alters the expression platform or the transcription process, known as riboswitches (Mandal and Breaker 2004; Winkler and Breaker 2005; Serganov and Patel 2007a). These studies involve a specified length of a noncoding region that undergoes conformational changes, and some of them have been studied at the transcriptional level as well (Heilman-Miller and Woodson 2003; Wickiser et al. 2005; Wong et al. 2007).

Transcription is a directional (5’-3’) process that involves the folding of the RNA molecule as it emerges — this event known as the sequential folding or co-transcriptional folding of RNA (Lai et al. 2013). Within recent years, it became evident that cofold structures form and influence the functionality of RNA (Pan and Sosnick 2006; Zemora and Waldsich 2010). Within the cellular environment, nascent RNA folds at a similar time scale as the process of transcription occurs (Brehm and Cech 1983; Pan and Sosnick 2006), and the geometry of RNA molecules determines the functionality of RNA.

Since folding starts at the 5’ end of nascent RNA and 3’ end is involved after the completion of transcript, it leads to the formation of transient structures. These structures may have a different functional role during co-transcriptional folding. Therefore, precise folding and functioning of these structures are dependent on the speed of transcription. Flavin mono-nucleotide (FMN) riboswitch is the best example to understand the importance of co-fold structure formation during transcription (Wickiser et al. 2005; Zemora and Waldsich 2010). These structures are functionally active during the natural speed (50nt/sec) of transcription, and varying the transcription speed could result in the functionally inactive transcript. This implies that some of the cofold structures are transient and perform distinct functional roles that may not be present in the intact purified transcript (Liu et al. 2016; Incarnato et al. 2017).

Transcriptional pausing is critical in precise co-transcriptional folding and is more likely to occur with specific pausing sites (Gabizon et al. 2018). Therefore, pausing could also be a significant mechanism that allows the transient structure formation and may be present for a specific duration only. It’s important to consider that pausing at specific sites could influence nascent RNA structure in a significant manner (Wickiser et al. 2005; Wong et al. 2005; Wong et al. 2007). Transient structures may promote specific functions by promoting conformational rearrangement (Perdrizet et al. 2012).

In this study, we identified the noncoding region of *ppgk* gene of *M. tuberculosis* that plays a critical role in its expression. We revealed that non-coding region becomes cleaved during the transcription (Figure 1). We determined the cleavage site that resides at the immediate upstream of Shine -Dalgarno sequence. This cleavage activity remains in a positive correlation with the efficient translation of PPGK protein. Results of *in vitro* assays show that cleavage occurs at a low to moderate rate of transcription. The nascent transcript of *ppgk* noncoding counterpart has an intrinsic functional property that is mediated by the formation of local secondary structures. Also, mutational studies enabled to determine the significance of this cleavage activity for efficient expression of PPGK protein. This study allows an understanding of co-transcriptional folding and intricate functioning that is the integral mechanism of *M. tuberculosis* H37Rv.

**Figure 1.**
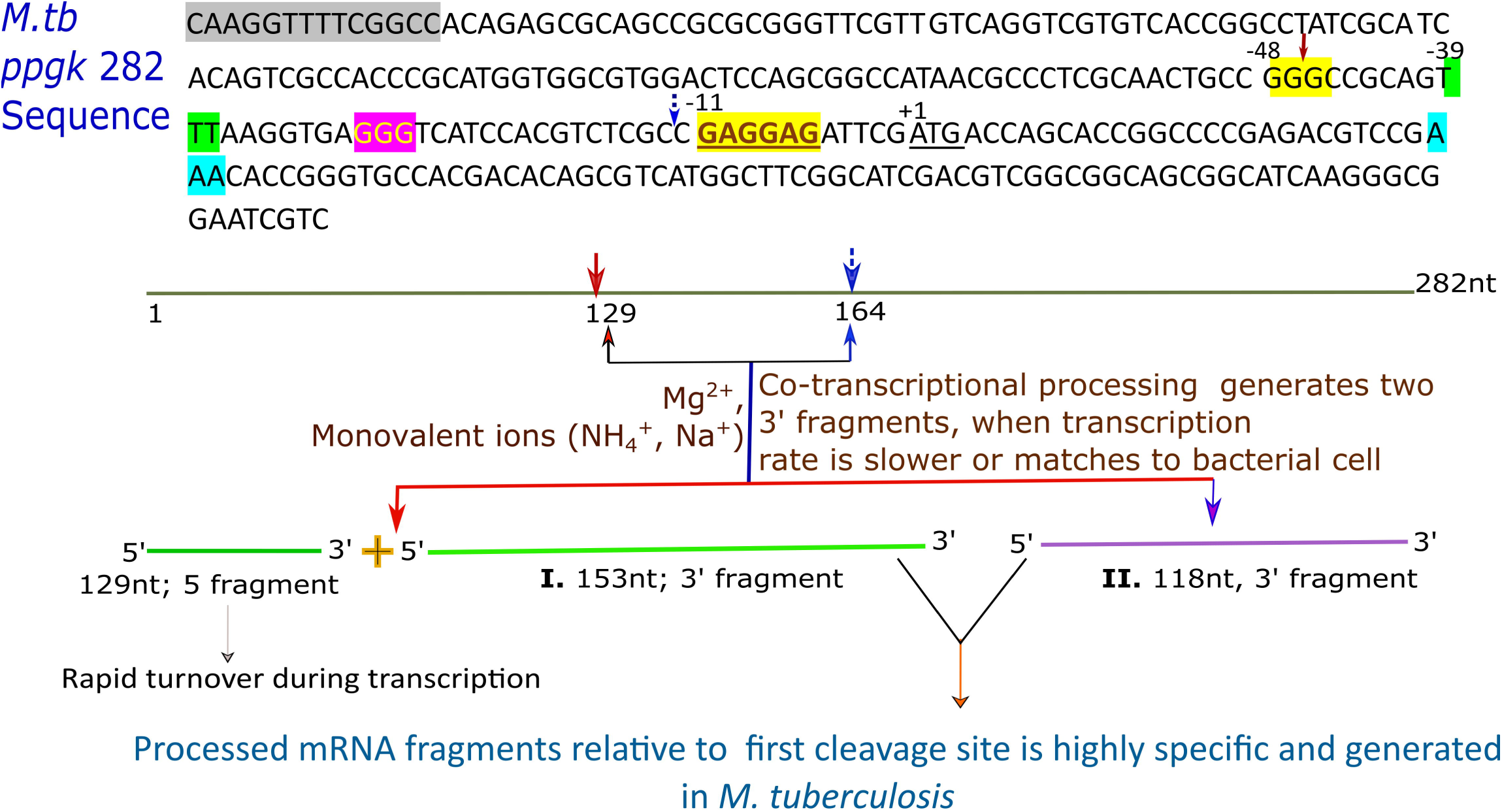
Overview of co-transcriptional mechanism mediated through 282-*ppgk* RNA element. Sequence represents the full length noncoding region, and 105 nucleotides of coding region. Highlighted nucleotides of the sequence have been used in the mutational study that are follows: grey: D1 (an initial stretch of TTTTCGGCC sequence); yellow: GGG; green: TTT; pink: GGG2; turquoise: AAA. Notation is given for the primary cleavage site with red arrow that generates 153 nt fragment, whereas second cleavage site is designated by blue arrow. The 282-*ppgk* RNA element undergoes simultaneous processing (two cleavage activities) during the transcription, and in the result two fragments generated that are dependent on the rate of transcription. Co-transcriptional processing is influenced by monovalent and divalent ions that determines ionic strength. Co-transcriptional processing of noncoding region occurs within *M. tuberculosis* at the same site, and mRNA fragments relative to specific cleavage site were precisely analysed by quantitative real time PCR.

## Results

### 5’UTR of *ppgk* gene is conserved among Mycobacterium species

The initial objective of this study was to identify a noncoding element that could have a potential role in the regulation of gene expression. We performed a sequence similarity search on several non-coding regions involved in the survival and virulence of *M. tuberculosis*. We found that iron dependent repressor (*ideR*) and polyphosphate glucokinase (*ppgk*) genes shared distinct similarities.

Further, we predicted secondary structures with the help of available web servers. The prediction was based on finding the probability for either Shine-Dalgarno sequestering domain or a hairpin loop formation that causes premature termination of a transcript. Although the non-coding region of *ideR* gene has shown high conservation among several Mycobacterium species, however, we could not deduce significant information from secondary structure prediction analysis of *ideR* noncoding element. On the other side, the noncoding counterpart of *ppgk* gene has shown considerable features that indicated the probability of gene regulation through a riboswitch or any specific molecular mechanism.

The full-length promoter of *ppgk* gene with the starting nucleotides position from −171 has already been reported. This full-length element (as a promoter) is also required for the expression of *ppgk* gene (Hsieh et al. 1996). We extracted non-coding regions of *ppgk* gene from several Mycobacterium species, from whom this gene sequence has been predicted or characterized as polyphosphate glucokinase. These regions were subjected to multiple sequence alignment by using MUSCLE software (Edgar 2004), to identify a consensus region. The noncoding region of *ppgk* gene starting from −165 is highly conserved among several species of Mycobacterium (Figure 2). Specific features of the noncoding element are the pseudoknot interaction stretch, cleavage sites, and SD sequestering region that remains consensus within the alignment. Accordingly, we selected the region spanning from −177 to +105 (282 bp) for further study. We considered the extended coding region (+105 relative to translation start site) as the adequate expression platform that was fused “in frame” with the *lacZ* gene, where the effect on β-galactosidase expression was tested. Secondly, primer pair was appropriate within the noncoding and coding region for PCR and made further experimental analysis feasible (including illumina sequencing and PAGE experimentations).

**Figure 2:**
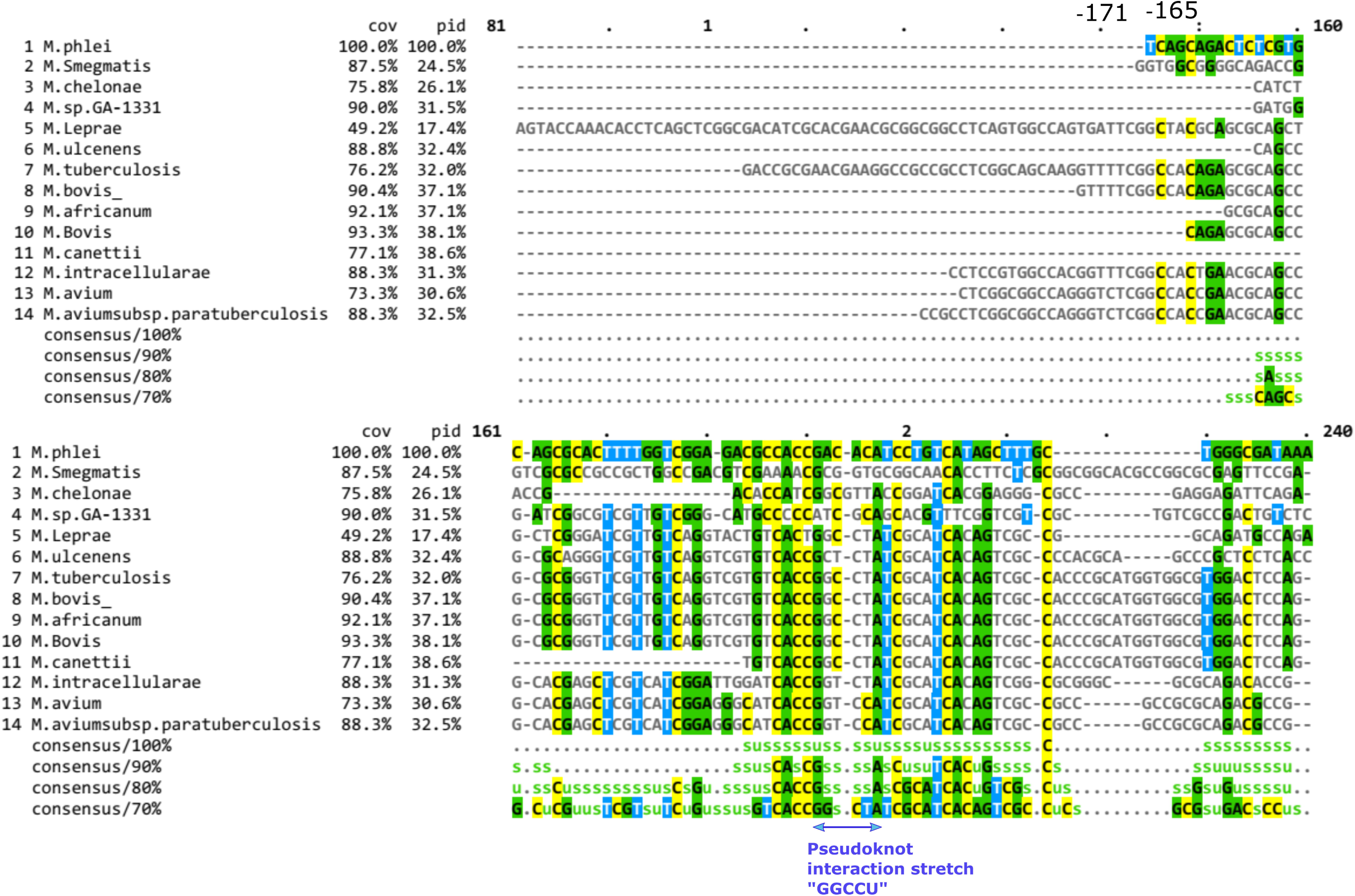
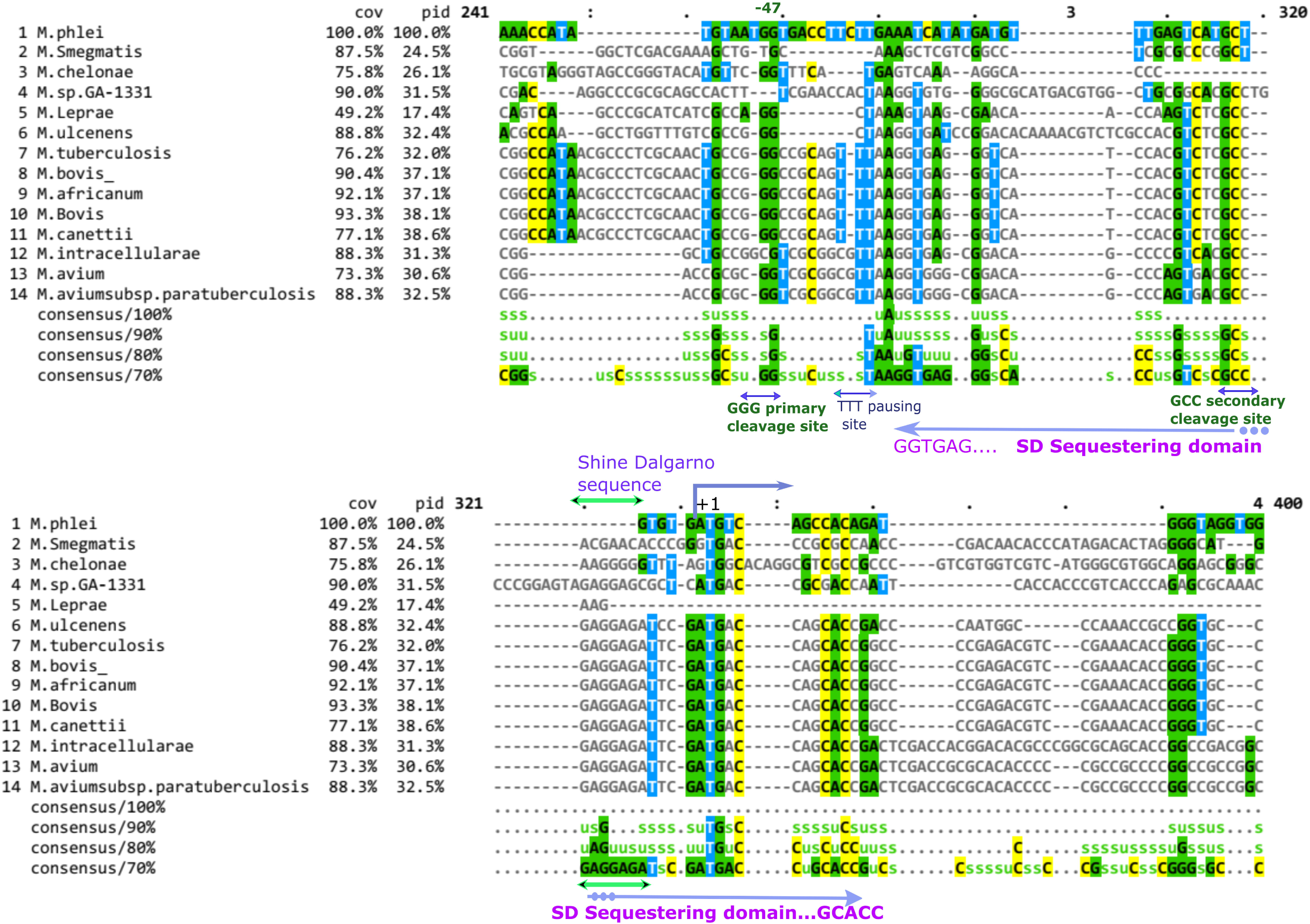
Multiple sequence alignment of non-coding region of *ppgk* gene. Noncoding regions were extracted from the complete genome of Mycobacterium species, from whom *ppgk* gene was either predicted or functionally characterized. These regions were aligned with the help of MUSCLE software that present the consensus nucleotides in different colour. Alignment shows that nucleotides position from −165 are conserved. Specific conserved features of the sequence alignment are pseudoknot interaction stretch (GGCCU), primary cleavage site (GGG) and secondary cleavage site (GCC). Region specific to upstream and downstream of SD sequence is conserved and features remain consensus within the alignment. The arrow shows the nucleotides region specific to SD sequestration.

The secondary structure prediction of the *ppgk* noncoding counterpart showed a specific hairpin region that sequesters Shine-Dalgarno sequence and initiation codon (predicted by Vienna RNA fold Server, Kinefold server, mfold web server including 3D structure by simRNA webserver as well) (Supplementary Figure 1A). The predicted probability for sequestering domain formation was >90%. This feature implied the possibility of regulation of *ppgk* gene that could be mediated by riboswitch or any specific mechanism. We designated this fragment as 282-*ppgk* RNA element in this study.

### *In vitro* transcription and illumina sequencing identified the cleaved 3’RNA fragment of *ppgk* noncoding region

The initial observations of this study were based on the *in vitro* transcription assays to define the functional insights of 282-*ppgk* RNA element with the transcriptional rates. The very first intriguing observation of this study was the generation of multiple transcripts as a result of *in vitro* transcription of a specific size of PCR product (Supplementary Figure 2). This observation was consistent with *the in vitro* transcription assays by using any bacteriophage RNA polymerase (T3, T7 and SP6). We considered the possibility of the second fragment as a result of using a transcription buffer of pH 7.9 containing 6 mM Mg^2+^ ions (provided by the manufacturer). We, therefore, optimized transcription buffer composition (pH 7.5) and standardized the Mg^2+^ ions concentration (10 mM). Under this condition, we observed a single band as a predominant product of transcription, with a low intensity of the second fragment (Figure 3A). The sizes of these fragments were corresponding to 282 nt (product size) and >100 nt. Therefore, it was pertinent to determine the possible catalytic activity or termination of 282-*ppgk* RNA element, and subsequent analysis of generated fragments by sequencing (5’ or 3’ end).

**Figure 3.**
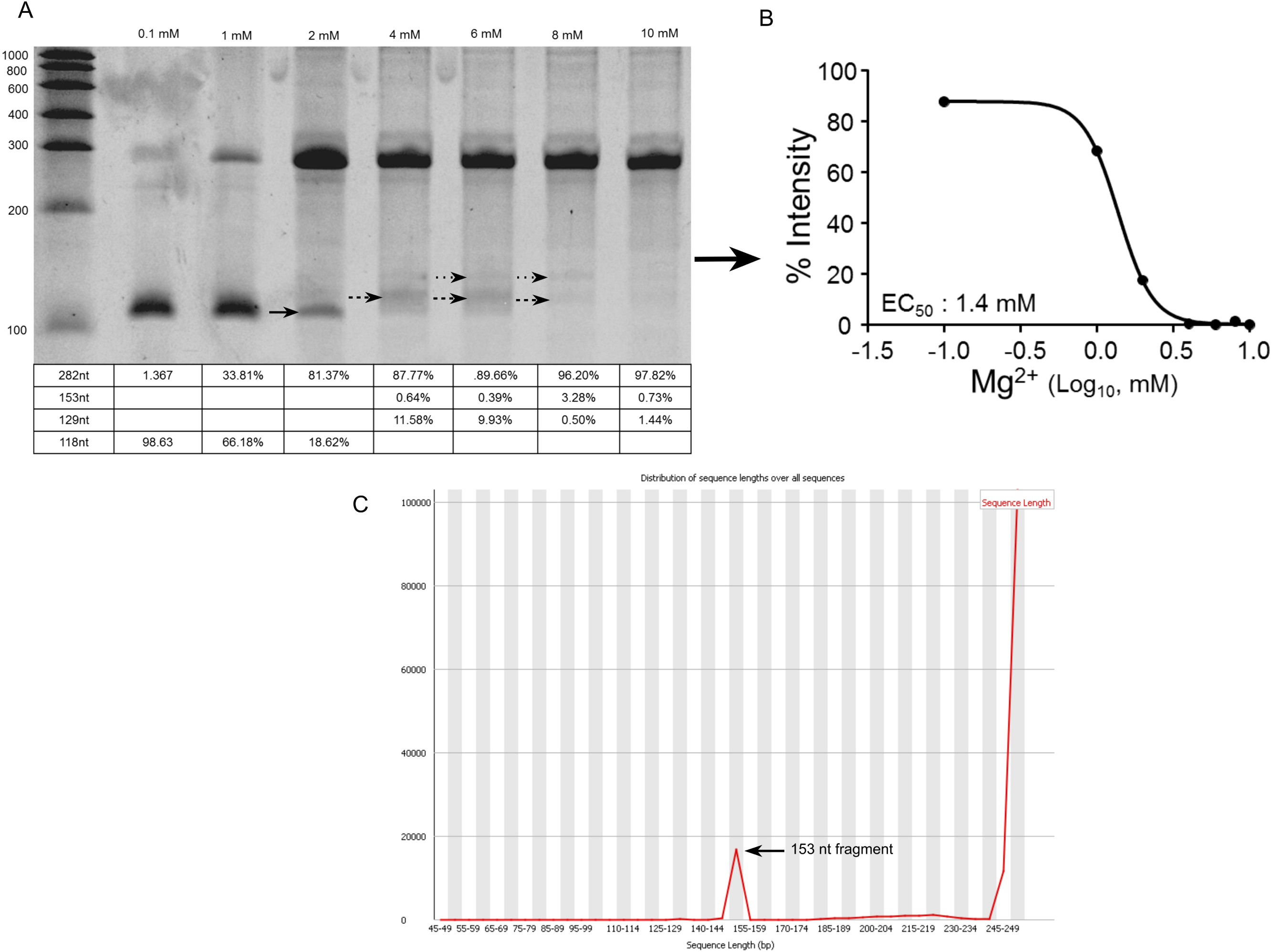
Co-transcriptional processing of 282-*ppgk* RNA element. (3A) *In vitro* transcription assay at different concentration of MgCl_2_ (0.1 mM-10 mM), and separation of transcripts on denaturing urea PAGE. Specific bands are shown with arrowhead, 153 nt (→); 129 nt (→) and 118 nt (→). Within the range of 0.1-2mM Mg^2+^ ions a single cleaved product (118 nt) observed, as the result of transcription. When the concentration of magnesium ions increased from 2-4 mM two distinct bands corresponding to 153 nt (3’ fragment) and 129 nt (5’ fragment; dispersed) can be observed. Moderate level of Mg^2+^ ion concentration (4 mM – 8 mM) maintains the transcription rate, and simultaneously correlated with the observed cleaved fragments. Percent intensity of each band is given as average from two biological replicates. Plot (3B) depicts the effect of Mg^2+^ ion concentration on the intensity of cleaved fragment (118nt and 153nt). Effective concentration_50_ (EC_50_) of magnesium ions (1.4mM), indicates that low rate of transcription is critical in maintaining the co-transcriptional processing. (3C) Graph is representative of sequence length distribution from the pattern file of 153 nt fragment that was revealed by illumina sequencing. This fragment contains SD-sequence and expression platform.

We performed an *in vitro* transcription assay by varying the concentration of Mg^2+^ ions. Since, Mg^2+^ ions are directly involved in the addition of NTP’s by coordination and alignment of phosphate groups (α, β and γ) (Nudler 2009), it would allow us to observe the transitional features of the 282-*ppgk* RNA element and help in the visualization of specific fragments that are generated with the simultaneous variation of transcription rate. To identify the 5’ or 3’ fragments, we transcribed the 282-*ppgk* RNA element at 4 mM and 8 mM Mg^2+^ ions and subjected to sequencing on illumina HiSeq2500 platform. These concentrations were selected on the basis of denaturing PAGE observations, where the possibility of 5’ and 3’ RNA fragment exists (Figure 3A).

RNA fragments were processed in integral form during library preparation, and sequencing reads were generated in the format of 2×100bp paired-end. Pattern files containing overrepresented sequences were used for analysis (Table 1). We analyzed the overrepresented pattern files that represent the complete 282 nt transcript, terminated transcript, or cleaved transcripts. Among the overrepresented sequences, only one sequence was determined to be specific for 3’ region of transcript that was further analysed from the corresponding pattern file (Figure 3C; Supplementary data file 3). This sequence started from the −47 positions (relative to initiation codon) and 36 nt upstream of the Shine-Dalgarno sequence. The length of this fragment was 153 bp and had shown a distinct peak in the plot of the distribution of sequence length (Figure 3C). However, we did not observe any pattern file corresponding to the terminated transcript.

During the variation of MgCl_2_ concentration in transcription buffer, we observed that at 0.1 mM to 2 mM concentration of Mg^2+^ ions, 118 nt fragment was of significant intensity, but during the transition from 2mM to 4mM and further, shifting in bands could be observed (Figure 3A). At 4mM and 6mM Mg^2+^ ions two new fragments were observed (5’ fragment 129nt; and 3’ fragment 153 nt), along with intact 282 nt fragment. The disperse region indicates the possibility of a rapid turnover of 5’ fragment during the co-transcriptional processing. These features imply the probability of undergoing two simultaneous cleavage activities within the nascent transcripts under the low rate of transcription. More specifically, 118 nt fragment is the result of two concurrent catalytic activities; thus, 153 nt fragment could not be a distinct product at a very low rate of transcription. The relationship between Mg^2+^ ion concentrations and cleaved fragment intensity is sigmoidal (Figure 3B).

### Non-coding RNA element of *ppgk* gene undergoes cleavage activity during the transcription

Since the transcription rate within the bacterial environment is much slower (20-80nt/sec) in comparison to *in vitro* (1000nt/sec), we performed *in vitro* transcription assay by varying the concentration of NTP’s and observed a similar pattern of bands (Figure 4). During the variation of NTP’s concentration within the 100 µM to 300 µM range revealed two major fragments. As the concentration of NTPs increased (400µM-600µM), concomitantly, the intensity of lower size fragment (∼100nt) decreased, and a new fragment (153nt) was observed. We determined the sequence of this specific fragment (∼100nt) by 5’RACE followed by sequencing. The sequence of this particular fragment (118 bases) starts precisely one nucleotide upstream from the SD sequence (C<u>GAGGAG</u>). In this regard, co-transcriptional processing was significant at the low rate of transcription (100-300µM NTP’s), whereas at the high rate of transcription, this activity was reduced (Figure 4). At 200 µM NTP’s, the observed intensity of 118 nt fragment was highest. A previously reported study has shown for FMN riboswitch in *Bacillus subtilis* that using 200 µM concentration of NTP’s results in a low level of full-length transcripts, in comparison to 500 µM NTPs. It indicates that co-transcriptional folding occurs at a lower rate of transcription, and FMN binding leads to termination of transcription (Wickiser et al. 2005).

**Figure 4.**
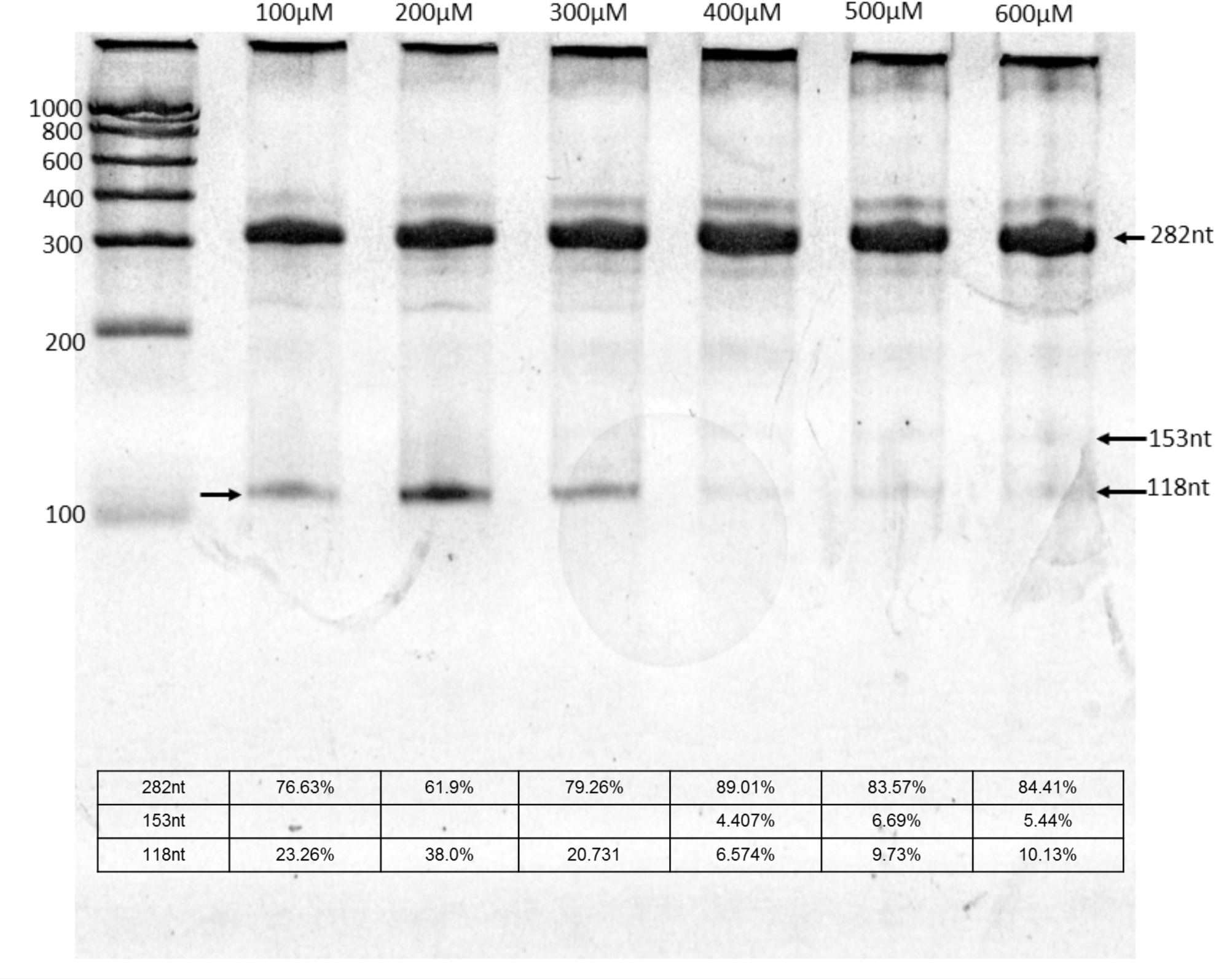
Concentration of NTP’s is correlated with co-transcriptional processing. At 100µM to 300 µM NTP’s intensity of 118 nt fragment ranges from 20% to 38%, that decreases to ∼9% at the optimum concentration (500µM) of NTP’s. NTP’s concentration within the range of 400 µM-600 µM, a different band of light intensity could be observed that is corresponding to 153 nt fragment. Provided band intensity is the average from two biological replicates.

### Cleavage activity within the non-coding region has a positive correlation with the expression of *ppgk* gene

RNA molecules have catalytic activity and also, additionally functioning in either rolling circle replication, transcription termination, expression or splicing (Cech et al. 1981; Buzayan et al. 1986; Teixeira et al. 2004; Winkler et al. 2004; Serganov and Patel 2007b). In this context, we sought to correlate the cleavage activity with the expression of PPGK protein. Since *M. smegmatis* and *M. tuberculosis* exhibit high sequence identity of RNA polymerase and transcription proteins, we, therefore, used *M. smegmatis* as a model in this study (Hubin et al. 2017). We introduced 5’UTR-*lacZ* constructs into the *M. smegmatis* mc^2^155, and analyzed the effect of 5’UTR on *lacZ* expression (Supplementary Figure 3). We analyzed specific mutations at the cleavage site (GGG) and possible pausing sites (TTT). Deletion mutant of three uracil residues leads to the abolishment of expression and has also shown significant reduction of cleaved fragments (Figure 5A). It leads to the conclusion that these uracil residues may be critical in maintaining the transient pausing and precise catalytic (cofold) structures during transcription.

**Figure 5.**
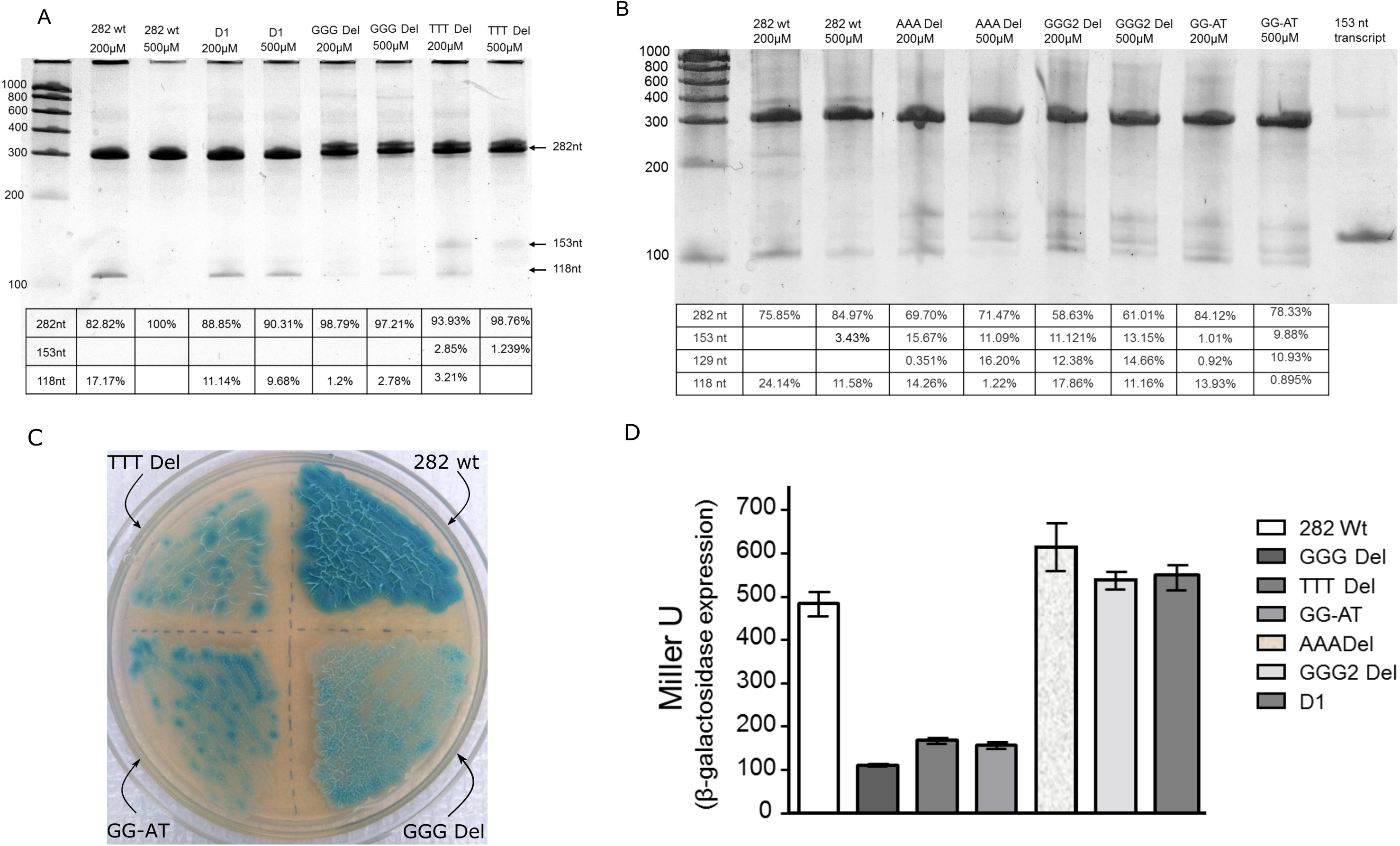
Effect of site specific mutations on co-transcriptional processing and reporter gene expression. Analysis of cleavage activity at 200 µM and 500 µM NTPs. (5A) *In vitro* assay of GGG and TTT deletion mutants have shown significant reduction in cleavage activity, and resultant 118 nt cleaved fragment. Transcripts of D1 mutant undergo cleavage activities at both low (200µM) and higher rate (500 µM NTP’s) of transcription. (5B) Effect of deletion mutants at the AAA and GGG2 sites on co-transcriptional processing. Transcripts of these mutants have shown fractional 153 nt and 129 nt bands. However, the effect of GG-AT mutation (at primary cleavage site) has shown reduction in cleavage activity and generation of 153 nt fragment at lower rate of transcription. 153nt fragment was transcribed and electrophoresed in parallel. Percent intensity of each band is mentioned in the respective lane of the table that is the average intensity from two biological replicates. (5C) Gene fragments carrying the site specific mutations within 282-*ppgk* element were cloned at the immediate upstream of the *lacZ* gene of pML163 plasmid, and transformed in to *M. smegmatis* mc^2^ 155 for β-galactosidase measurement. Transformed cultures were streaked on 7H10 plate, supplemented with 10% ADC and 20 mg/ml X-gal. (5D) Level of β-galactosidase expression from wild type and mutant samples. All of the samples were analysed from three biological replicates, and average miller units were plotted. Error bars indicate the standard deviations (mean values) from three biological replicates.

Deletion mutant at the cleavage site (GGG) leads to a reduction of co-transcriptional cleavage activity at 200 µM concentration of NTP’s (Figure 5A), and loss of β-galactosidase expression (Figure 5D). The level of β-galactosidase did not vary significantly when some initial stretch (TTTTCGGCC; D1 mutant) of noncoding counterpart was deleted. Thus, the correlation between cleavage activity and expression is of major significance. It is important to consider that variation in the intensity of any fragment (153 nt or 118 nt) provides a direct correlation with an expression where SD sequence has a critical role.

We analyzed the effect of mutations at other sites of the sequence as well. Analysis of transcription products of AAA and GGG2 deletion mutants did not exhibit a reduction in cleavage activity and thus neither affect the expression as well (Figure 5B and 5D). AAA deletion constructs increased the expression marginally, indicating that co-transcriptional folding is partially affected by the coding region and thus affect the processing at the same time as well. These sites are present at the downstream of the first cleavage site that generates 153 nt fragment; therefore, it did not affect the generation of 153 nt fragment. However, noteworthy to mention GG-AT mutation (at cleavage site) where the generation of 153nt fragment (and respective 5’ fragment of 129nt) was significantly reduced (1.01%) at low rate of transcription and abolished the expression as well (Figure 5B, 5D). Although, in this case, 118 nt fragment remains as the major cleaved fraction. The significant conclusion from all of these observations is that simultaneous processing occurs under *in vitro* condition, where the generation of either 3’ fragment is correlated with protein expression.

### Evaluation of cleaved *vs* intact fragment ratio of *ppgk* transcripts in the *M. tuberculosis* H37Rv

Transcription within the bacterial cell occurs at the 20-80nt/second, which is much slower in comparison to *in vitro* studies (Heilman-Miller and Woodson 2003); therefore, the nascent transcript of *ppgk* noncoding region might have effective and critical functioning in *M. tuberculosis*. Within the perspective, we analyzed the levels of *ppgk* transcripts that are either in full length or cleaved within the bacterial cell. Purified RNA from *M. tuberculosis* was subjected to cDNA synthesis by using *ppgk* gene-specific primer. We examined the existence of transcripts that retained the noncoding region or cleaved fractions. The copy number of a cleaved fragment (*ppgk* mRNA I) exceeded by 1.69 fold in comparison to the intact transcript (containing extended noncoding counterpart) (Figure 6). The copy number of *ppgk* mRNA relative to the second cleavage site (*ppgk* mRNA II) had nearly equal Cq value with a fold change of 1.58 (Supplementary Figure 3).

**Figure 6.**
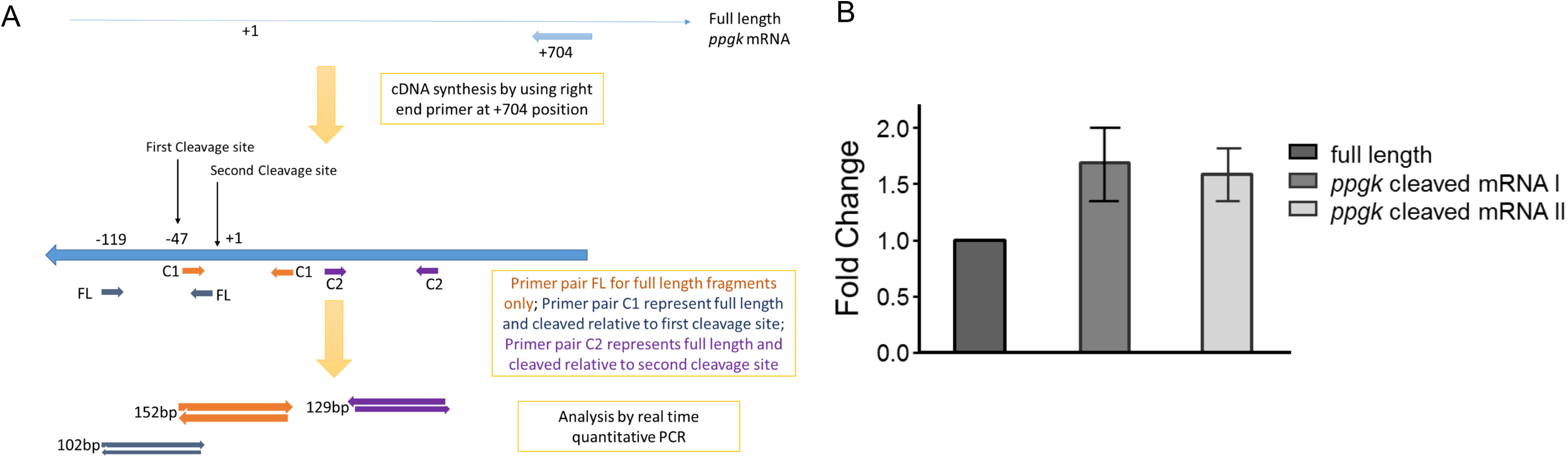
Analysis of cleaved *vs* intact fragment ratio of *ppgk* mRNA level within *M. tuberculosis*. (A) Reverse transcription was performed by using right end *ppgk* gene specific primer, followed by alignment of primer pairs those were used in the real time quantitative PCR. These primers were used respectively according to specific cleavage site, and therefore quantified fractions would be evaluated as intact *vs* processed transcripts of *ppgk* gene in *M. tuberculosis*. It could be noted that left primer for full length fragment analysis would anneal with full length cDNA fragments only, and thus Cq value corresponding to complete fragments. Whereas, primer set for cleaved fragment would bind to full length cDNA and cleaved fractions as well (left primer starts with cleavage site). (B) Fold change values were determined from the absolute quantification. Cq values of full length transcripts were normalized to 1 and data obtained were plotted in fold change. Data were obtained from three replicates, average Cq values were determined and fold change values plotted. The cleaved mRNA fractions relative to first (*ppgk* mRNAI) and second cleavage site (*ppgk* mRNAII) have nearly equal fold change values, that is 1.69 and 1.583 respectively, revealing that *ppgk* mRNA undergoes cleavage activity (36nt upstream from SD sequence) during the transcription, and these mRNA fractions remain stable for expression. Results were presented as mean ± SD from biological replicates in triplicates.

In conclusion, we found that co-transcriptional folding and processing of *ppgk* transcript occurs within the *M. tuberculosis*. Importantly, processing of non-coding regions is a critical event in maintaining the cleaved transcripts; those, in turn, provide a stable expression platform for PPGK protein expression. These results also revealed that within *M. tuberculosis*, local secondary structures of nascent transcripts are critical in determining the functional roles.

### Monovalent ions have a predominant functional role in co-transcriptional processing

In this work, we investigated the involvement of two specific sites during the co-transcriptional processing of the noncoding region of *ppgk* gene. This molecular mechanism is highly sequence-specific and generates a single 3’ RNA fragment that is dependent on the transcription rate. The first reported ribozyme of 26S rRNA of *Tetrahymena* undergoes splicing activity in the presence of monovalent cation, divalent cation and GTP (Cech et al. 1981). In this process, the first step involves the phosphoester transfer reaction between the 3’-OH group of GTP and 5’-phosphate of the splice site. Subsequently, the second step involves the process of ligation and removal of intron.

Therefore, to understand the specific role of any of the element in the processing of non-coding region of *ppgk* gene, we performed *in vitro* transcription of 282-*ppgk* element and observed the increase in the intensity of 118 nt RNA fragment with the increment of ammonium sulfate concentration in transcription buffer (Figure 7A). The intensity of cleaved RNA fragment (118nt) increased, thus it could be speculated that monovalent ions could play a significant role in the co-transcriptional processing. Subsequently, we determined that the purified 282-*ppgk* RNA element has intrinsic activity in the presence of monovalent ions only (Figure 7B), and the requirement of Mg^2+^ ions could have a synergistic effect but not required for the generation of 153nt fragment (first cleavage site).

**Figure 7.**
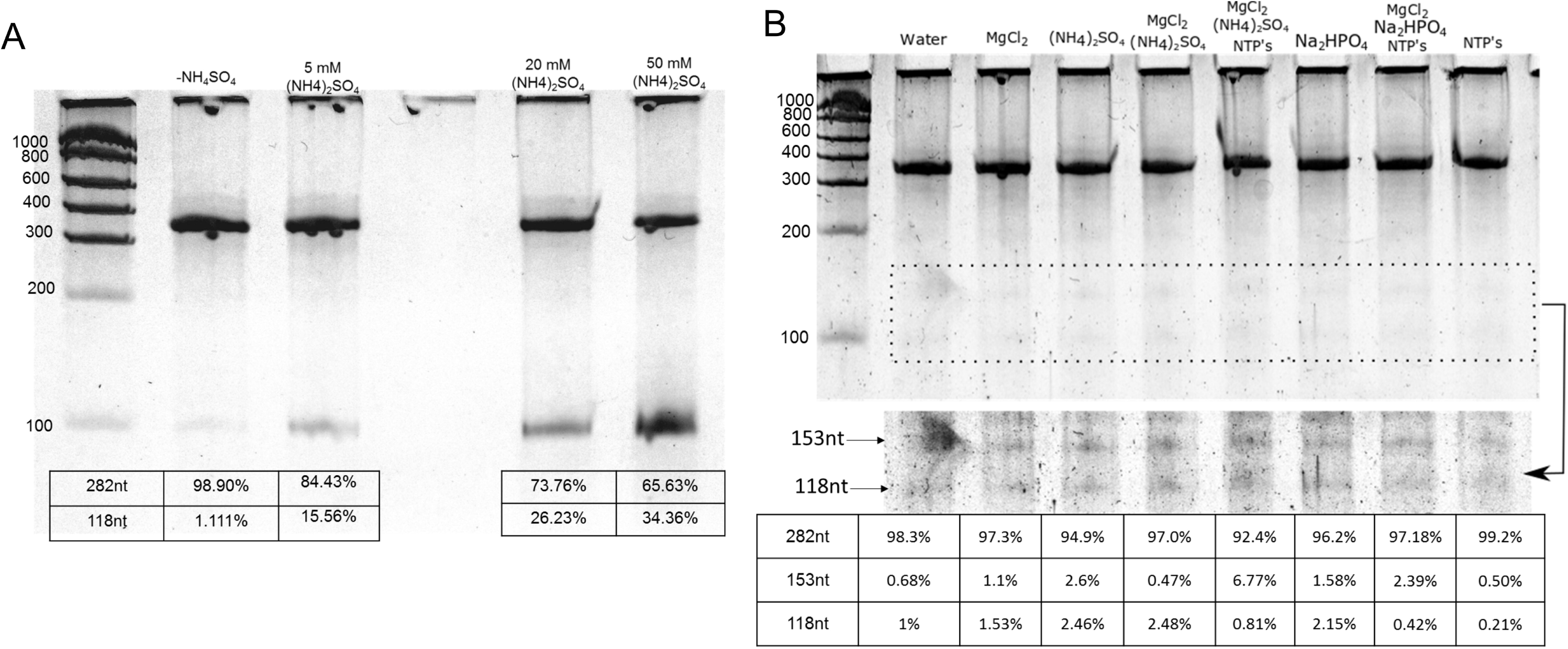
Co-transcriptional processing influenced by ionic strength in transcription buffer. (A)Transcription assay was performed by increasing ammonium sulphate concentration and keeping the constant MgCl_2_ concentration (6mM) in transcription buffer. Intensity of 118 nt fragment increases with the presence of salt ions in transcription buffer. (B) Intrinsic activity of purified 282*-ppgk* RNA element influenced by salt ions (15 mM). Purified RNA undergoes independent cleavage activity in the presence of salt ions. Selected region (rectangular) was enhanced by increasing white balance with brightness and contrast adjustment, although complete gel image was used for analysis (without enhancement). Noteworthy to mention about the 153 nt fragment, that has increased intensity in the presence of 15mM (NH_4_)_2_SO_4_ only. Although, the intensity of band is slightly higher when incubated in combination with 5 mM MgCl_2_. Effect of Na_2_HPO_4_ remains equal or in combination with MgCl_2_ and NTP’s. Concentration used in the experiment-ammonium sulphate [(NH_4_)_2_SO_4_] at 15 mM, disodium hydrogen phosphate [Na_2_HPO_4_] at 15mM, MgCl_2_ at 5 mM and NTP’s at 100 µM. Samples incubated at 37°C for 40 minutes.

## Discussion

### Secondary and tertiary structure determines the fate of ongoing nascent transcript

Most of the riboswitches are controlled either at the level of transcription termination or translation inhibition. Regulation of translation mediated by the formation of Shine-Dalgarno sequestering domain that prevents the access of ribosome binding (Nahvi et al. 2002; Winkler et al. 2002; Serganov and Patel 2007b) is the major feature of some of the discovered riboswitches. In this work, we identified a novel non-coding element that has predominant intrinsic activity at low to moderate rate of transcription, where a low range of magnesium ion concentration (1-2 mM) is significant in co-transcriptional processing. The noncoding counterpart of *ppgk* gene has shown distinct secondary structure, including sequestration of Shine-Dalgarno sequence and initiation codon. We strongly propose that the formation of SD sequestering domain is the underlying mechanism that does not allow PPGK expression from an intact transcript (including full-length noncoding region). This mechanism was supported by site-specific deletion mutants, where cleavage activity also affected the reporter gene expression. Prediction and analysis of three-dimensional structure show the possibility of “in-line” nucleophilic attack that could be the basis within the “GGG-turn” (with 28.3° angle) at the cleavage site (Figure 8A). Three or two guanine nucleotides are conserved at the same position among several species of Mycobacterium, indicates that such activity might occur among these species as well (Figure 2). An important aspect among these species is the presence of SD sequence and other important conserved elements, for example, primary cleavage site and secondary cleavage site as well. Apart from this, pseudoknot interaction stretch makes an important contribution to the functioning of the noncoding element.

**Figure 8.**
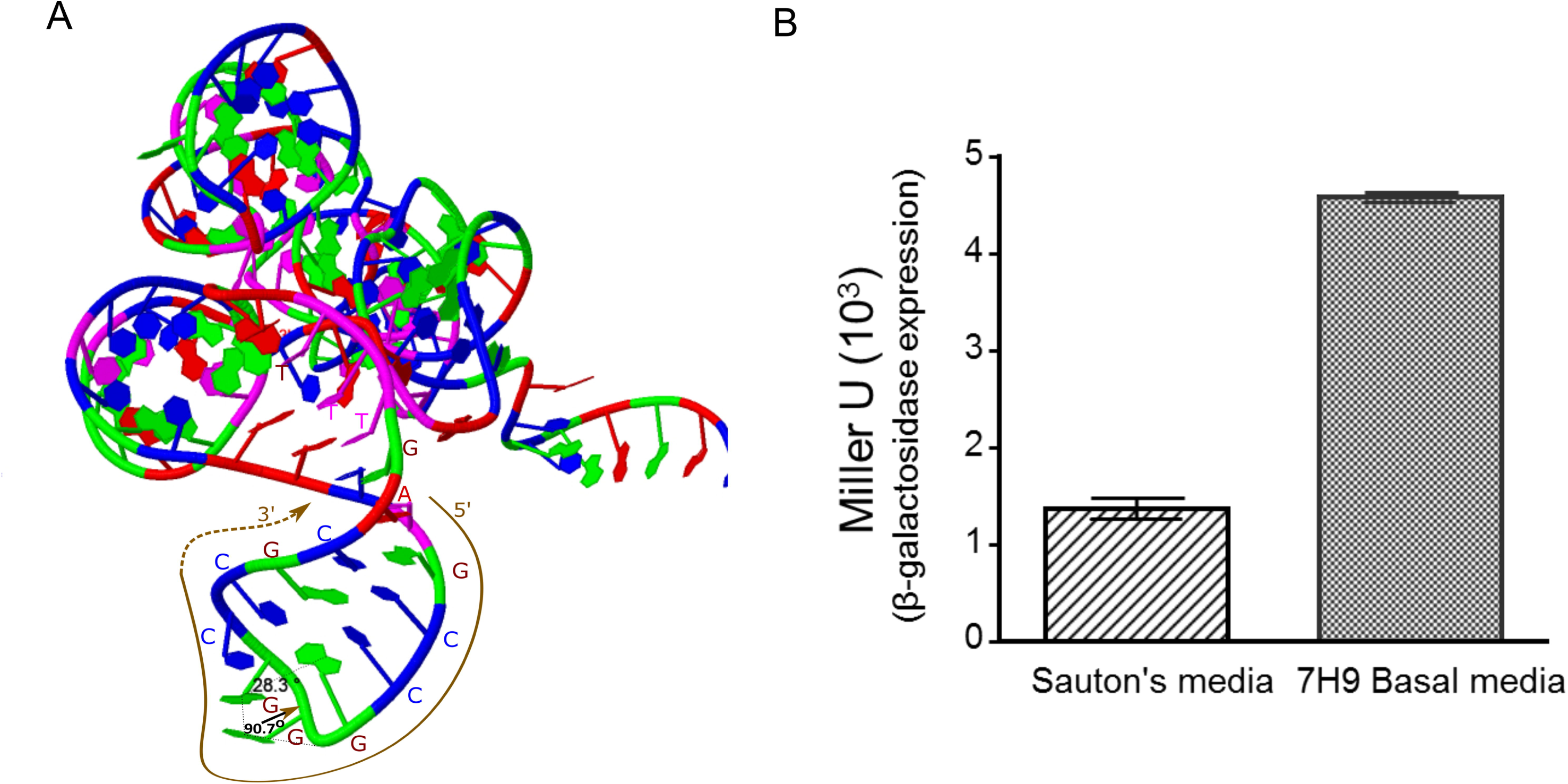
(A) Three dimensional Cofold structure of a specific noncoding region (−162 to −23) of 282-*ppgk* RNA element, analysed by simRNA webserver. Input sequence for 3D structure determination was specified with secondary structure notations that was derived by cofold server (for same sequence), thus it allowed to observe 3D cofold structure within the context of cleavage site. PDB file visualized by Jena 3D viewer and angles were measured between specific nucleotides. Three consecutive guanine nucleotides (in red) makes a sharp turn (28.3°) that follows cleavage site (shown by arrow). (B) Analysis of β-galactosidase expression when cells were allowed to survive in basal Sauton’s broth medium, and 7H9 medium salts (without any supplement). Although, O.D._600_ was not changed under both medium (remains ∼0.2 after three days), nonetheless reporter gene expression was significantly higher in 7H9 salt based medium because of the higher amount of salts composition and other divalent ions. Error bars represent the standard deviations from three biological replicates.

Additionally, we observed that the ionic environment and concentration of Mg^2+^ ions could play a significant role within the Mycobacterium, where the non-coding region remains in a positive correlation with reporter gene expression. β-galactosidase expression slightly increased when cells were grown in the presence of MgCl_2_ (Supplementary Figure 5). In a different experiment, cells were allowed to survive in the presence of basal Middlebrook 7H9 medium and Sauton’s broth without the addition of carbon source (glycerol or ADC enrichment). The supplement was not added to exclude the role of a metabolite that may influence the expression or to determine the reporter gene expression in the presence of salts only. Though O.D. _600_ did not vary after three days, however expression was significantly increased in the presence of 7H9 medium in comparison to Sauton’s broth medium (Figure 8B). This could be because of co-transcriptional folding that is influenced by the ionic environment and consequently processing as well, as we observed the cleaved fragments in the presence of monovalent ions only (Figure 8B).

Importantly, folding kinetics of RNA secondary structures is prominently influenced by salt concentrations, though Mg^2+^ ions have a comparatively stronger effect (Hanna and Doudna 2000; Pan and Sosnick 2006; Li et al. 2008). It is important to mention that the composition of 7H9 medium is rich in ammonium sulfate and disodium hydrogen phosphate along with other salts. Therefore, the ionic environment could alone be a critical determinant in maintaining the efficient processing of nascent RNA and its expression.

### Nascent transcript of *ppgk* noncoding region has predominant catalytic activity

Here we identified the noncoding RNA element that undergoes co-transcriptional processing during the low to moderate rate of RNA synthesis that also describes the significance of this event within the bacterial cell. Specific cofold structures of nascent RNA might be transient and rearranges within the intact transcripts. We observed an increase in the intensity of cleaved fragment after incubation of 282-*ppgk* RNA element in the presence of increasing concentrations of Mg^2+^ ions (Figure 9). The crucial observation was to visualize the fragment intensity of 129 nt and 118 nt, and both have increased up to 5.06% and 6.15%, respectively (at 10 mM MgCl_2_). However, at the same time, only a fractional increase (2.97%) in the intensity of 153 nt fragment was observed.

**Figure 9.**
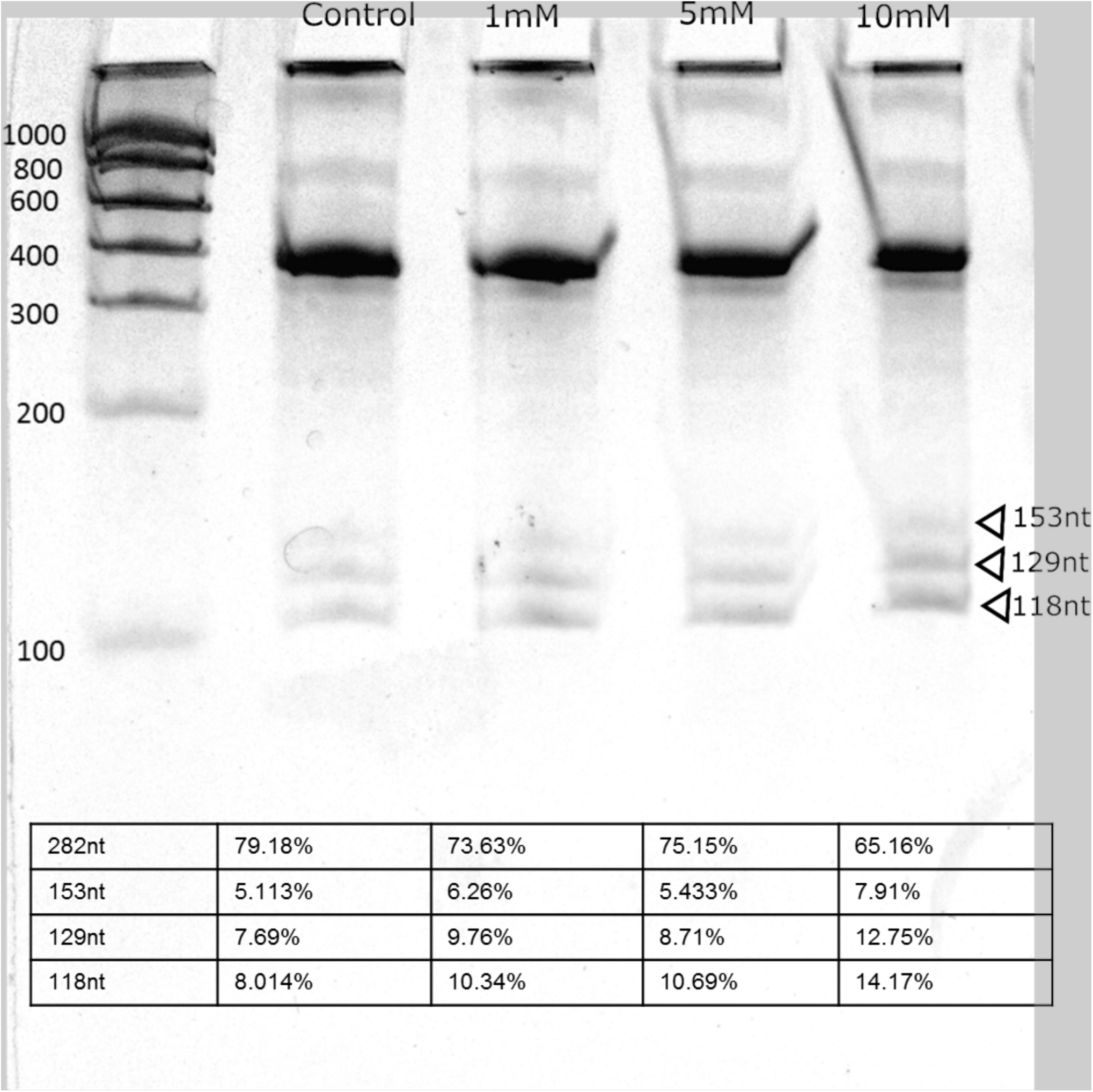
Analysis of cleavage activity of 282- *ppgk* RNA element under protein free environment. Transcribed RNA was purified by phenol; chloroform extraction and incubated at different concentrations of Mg^2+^ ions. We analysed the products on denaturing PAGE and followed by silver staining. As the concentration of Mg^2+^ ions increases concomitantly 129 nt and 118 nt fragment intensity increases, but the intensity of 153 nt fragment does not increase significantly.

**Figure 10.**
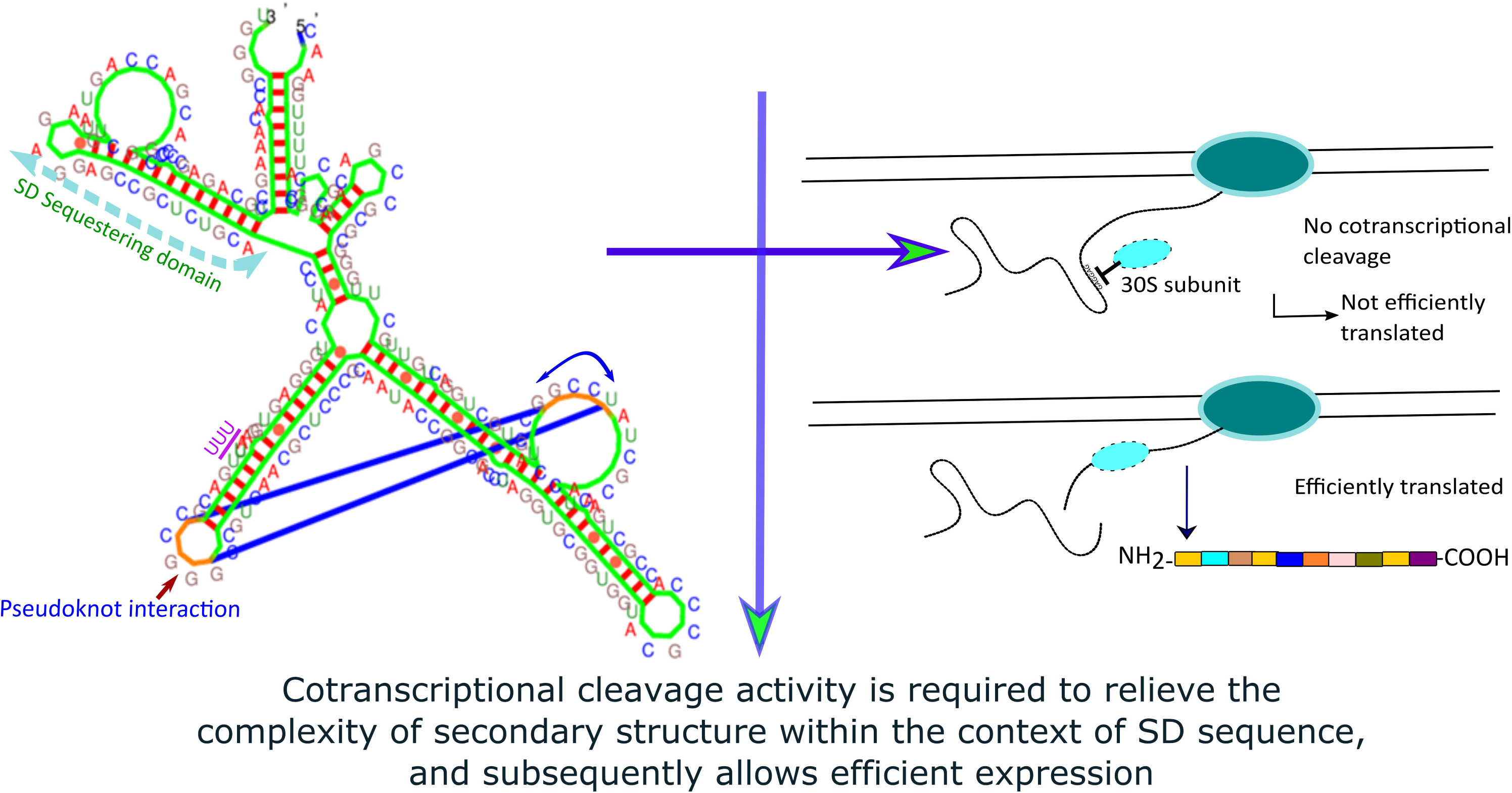
Conclusion of co-transcriptional processing mediated by non-coding RNA element of *ppgk* gene. During the transcription, non-coding RNA fragment becomes cleaved at the upstream of ribosomal binding site. This event is required for relieving the secondary structure of mRNA and efficient binding of ribosomal subunit for expression. Full length transcript would be insignificant in the expression of protein because of the sequestration of ribosomal binding site.

It does allow us to conclude that both cleavage activities co-occur within 282-*ppgk* RNA element, and thus the intensity of 153 nt fragment was not increased in a significant manner. Besides, it is significant to consider that primary cleavage activity that is in correlation with 153 nt fragment generation might be an intrinsic property of nascent RNA and plays a predominant role in expression as well. We did not visualize any particular catalytic activity of PAGE purified 282-*ppgk* RNA element upon post-incubation in the presence of divalent metal ions (Supplementary Figure 6). The possible explanation is that there are critical secondary structural features (pseudoknot interactions; supplementary figure 7) that remain stable for the specific time duration (or transiently stable), and could not be restored once denatured.

Overall experimental findings indicate that 282-*ppgk* RNA element has active cofold intermediates (secondary structures) during transcription and undergoing processing at the same time. Although this study was done under *in vitro* condition, nevertheless, we verified the existence of cleaved transcripts in *M. tuberculosis* with respect to the same cleavage site. Moreover, cleaved transcripts are higher in *M. tuberculosis* in comparison to intact transcript (carrying full-length noncoding region). It provides the significant possibility of cleaved transcripts in maintaining the stable and efficient expression of PPGK enzyme.

### Noncoding RNA element of *ppgk* gene could not be a riboswitch

Experimental results have shown that the non-coding RNA element of *ppgk* gene could not be a riboswitch element. There are two possible explanations; first is the observation of β-galactosidase expression in the presence of salt-based medium (7H9 and Sauton’s broth) only. Secondly, we did not observe a significant difference in reporter gene expression in the presence of specific divalent metal ions or metabolites (glucose and glucose-6 phosphate) (Supplementary Figure 5). It directly indicates that the genetic expression of *ppgk* gene is a constitutive process in *M. tuberculosis*, where the co-transcriptional processing is required to alleviate and maintain the ribosomal binding site for efficient expression.

## Methods

### Oligonucleotides, chemicals, and enzymes

Oligonucleotides were purchased from Integrated DNA Technology (IDT) (Supplementary Table 1). Custom gBlocks gene fragments were purchased from Integrated DNA Technology. All of the enzymes were purchased from New England Biolabs (NEB), and chemicals were purchased from Sigma. Ultrapure urea was purchased from Affymetrix Invitrogen.

### Programs and Software used for sequence alignment and structure determination

Specific noncoding region of *ppgk* gene from *M. tuberculosis* was subjected to BLAST analysis using Refseq Representative Genome Database. Multiple sequence alignment was done for intergenic sequences of *ppgk* gene by MUSCLE (MUltiple Sequence Comparison by Log-Expectation) software. This software provides high accuracy and precision within the alignment and enables to determine critical residues.

Co-transcriptional folding path of 282-*ppgk* RNA was simulated by using kinefold webserver that is based on the exactly clustered stochastic (ECS) algorithm. Prediction and analysis of secondary structure were performed by using Vienna RNA fold Server, Kinefold server, and mfold web server (Hofacker 2003; Xayaphoummine et al. 2005). Cofold secondary structure was predicted by using Cofold webserver (Proctor and Meyer 2013), and output secondary structure sequence was used for 3D cofold structure prediction. Three-dimensional structure was predicted by using simRNA webserver and PDB file was analyzed by Jena 3D viewer (Boniecki et al. 2015). All graphs were plotted using Prism 6 software.

### PCR amplification and *in vitro* transcription of a noncoding element of *ppgk* gene

Total genomic DNA from *M. tuberculosis* H37Rv was used as a template for PCR. Specific primers were used to amplify the region spanning from −177 to +105 (relative to initiation codon) of *ppgk* gene. T7 RNA polymerase promoter (<u>TAATACGACTCACTATA</u>GGG) was attached to the forward primer. Phusion high fidelity DNA polymerase kit (NEB) was used for the PCR amplification and optimized with DMSO and formamide. Confirmation of single PCR product was done on 2% agarose gel electrophoresis and native PAGE as well. PCR products were purified by QIAGEN Quick extraction purification kit, and eluted in RNase free MQ water. *In vitro* transcription was performed in 20µl final volume containing 40mM Tris-Cl (pH7.5), 2mM spermidine, 10mM NaCl, 10mM MgCl_2_, 10mM DTT, 800ng of PCR product and 20 units of T7 RNA polymerase. Transcription reaction was incubated at 37°C for one hour. Transcripts were purified by phenol: chloroform: isoamylalcohol (125:25:1) and followed by chloroform: isoamylalcohol (24:1) extraction. During extraction samples were centrifuged at 12,000g for 2 minutes and aqueous layer was precipitated by the addition of 0.5 volume of ammonium acetate (7.5M) and 2.5 volume of ethanol. Purified RNA was dissolved in RNAse free water.

### Sample preparation and sequencing by illumina HiSeq 2500 of 282-*ppgk* RNA samples

*In vitro* transcription of 282 nt fragment was performed in the presence of 4mM and 8mM MgCl_2_ containing transcription buffer. After one hour of incubation at 37°C, samples were purified as described above and dissolved in RNAse free water. In order to maintain the integrity of 5’ or 3’ RNA fragments, samples were not further processed for Ribozero or any fragmentation steps. Instead, RNA samples were directly processed for cDNA synthesis using standard protocol for library preparation and sequencing according to manufactures instructions.

One microgram of RNA was used for the library preparation by using TruSeq RNA Sample Prep v2 kit. Library was validated by using Agilent 2100 Bioanalyzer. Library was pooled and normalized.

Sequencing was generated on HiSeq 2500 system in the format of 2×100bp paired end. With the help of in house python script, overrepresented sequences were extracted in the form of pattern files. Each pattern file was separately analyzed.

### *In vitro* Transcription of 282-*ppgk* RNA and analysis of cleavage activity

*In vitro* transcription was performed as described above by varying the concentration of magnesium ions (0.1 to 10 mM) or NTP’s (100 to 600µM). During the variation of NTP’s concentration, we used transcription buffer of 8mM MgCl_2._ Transcripts were purified and dissolved in RNAse free water. Transcribed samples (1µg) were mixed with urea (18M; 2X) containing loading dye (Regulski and Breaker 2008). Samples were loaded on 8M urea (denaturing) PAGE (8%), and electrophoresed in 1X TBE buffer (90mM Tris–HCl, 90 mM borate, and 1 mM EDTA). Bands were visualized by silver staining as described (Chevallet et al. 2006), and the image was captured by using white UV transilluminator screen. All of the gel images were converted to grayscale with the help of GNU Image Manipulation Program (GIMP), and slight brightness and contrast were adjusted (to complete gel image) without affecting visualization of bands. Analysis of bands intensity was done by using CLIQS 1D Gel Image Analysis Software. Average bands intensity was calculated from the two biological replicates and provided with gel images.

To investigate the role of monovalent salt ions in co-transcriptional processing, we performed the transcription at different concentrations of ammonium sulfate, keeping the constant MgCl_2_ concentration (6mM in transcription buffer). For post-incubation analysis, transcribed RNA was purified, and 850ng was incubated at 37°C for 40 minutes, in a combination of salts, metal ions or NTP’s as mentioned. Ammonium sulfate and disodium hydrogen phosphate were used at 15mM, MgCl_2_ at 5mM and NTP’s at 100 µM concentration. Ammonium sulfate and disodium hydrogen phosphate are enriched components of Middlebrook 7H9 Broth.

### 5’RACE for co-transcriptional cleaved fragment sequencing

*In vitro* transcription of 282-*ppgk* RNA element was performed in the transcription buffer of 8mM Mg^2+^ ions in the presence of 200 µM of NTP’s. Samples were electrophoresed on denaturing (8M urea) PAGE, and bands were visualized under UV after ethidium bromide staining, bands specific to 118 nt fragment was eluted by crush and soak method (Regulski and Breaker 2008). The 5’ Phosphate group of the purified RNA was removed by Calf Intenstinal Phosphatase. The 5’-phosphorylated RNA was ligated to 5’ RACE adapter by using T4 RNA ligase. The ligated product was reverse transcribed by AMV RT into cDNA by using the right end gene-specific primer (Table 1). PCR product was amplified by 5’ adapter specific primer (outer) and 3’ gene-specific primer. PCR product was purified and sequenced by an adapter primer.

### Generation and analysis of specific mutants of noncoding element of *ppgk* gene

Sequences with desired alterations (Figure 1) were synthesized as gBLOCK gene fragments (carrying PmeI and Sph1 restriction sites). These fragments were directly cloned into the pML163 plasmid and transformed into *M. smegmatis* mc^2^155. Transformed colonies were selected on 7H10 agar (supplemented with 10%OADC) containing 50µg/ml hygromycin and 20mg/ml X-gal. Selected colonies were grown in 7H9 (supplemented with 10%ADC) media. Plasmids were isolated and sequenced to confirm in-frame fusion of 5’UTR with the *lacZ*.

To study the mutations in the fragments, gBLOCK gene fragments were amplified by PCR, and transcribed as described before. Transcripts were synthesized in the presence of 200 µM and 500 µM NTP’s, and 1µg of transcripts were analysed in parallel to wild type fragments.

### Transformation of 5’UTR-*lacZ* constructs and analysis of β-galactosidase expression within *M. smegmatis* mc^2^155

Total genomic DNA was isolated from *M. tuberculosis* H37Rv, according to a standardized protocol in the laboratory. The specific region from −177 to +105 (relative to initiation codon) was PCR amplified. PmeI and SphI digested PCR products were cloned into pML163 plasmid, at the immediate upstream of the *lacZ* gene that generates in-frame fusion with the initiation codon of β-galactosidase.

Transformed colonies of *M. smegmatis* mc^2^155 were grown in Sauton’s medium supplemented with glycerol, and 0.05% tween 80 was added to prevent clumping. To measure the LacZ expression in the presence of exogenous ligands, metabolites or metal ions were added when O.D. reached 0.2, and incubated further for ∼30 hrs. Cells were harvested at mid-exponential phase (O.D_600_ ∼ 0.8) and suspended in the same medium. Individual culture (three replicates) was sonicated by VibraCell Sonicator at 4°C. β-galactosidase measurement was done by using 400µl of a sonicated sample, mixed with the 600µl lacZ buffer (permeabilization buffer) and incubated at 28°C for 15 minutes. 200µl ONPG (4mg/ml) was added and samples were incubated at 28°C until yellow color is formed. Reactions were stopped by the addition of 500µl Na_2_CO_3_ (1M). Samples were centrifuged at high speed, and absorbance was measured at 420 and 550nm. Miller units were calculated as described (Malke 1993).

In a different experiment, to analyze the LacZ expression in the presence of 7H9 and sauton’s broth, the cultures were grown without any supplement (ADC, glycerol or tween). This was done purposefully to evaluate the β-galactosidase expression in the absence of metabolite. The transformed culture of *M. smegmatis* was allowed to survive in the presence of the ionic environment (salt ions) only.

### Total RNA isolation and real-time quantitative PCR

*M. tuberculosis* H37Rv, was cultured in 7H9 medium (supplemented with 10%ADC). After 12 days (mid exponential phase), culture was harvested, washed in TE buffer. The required amount of lysozyme and proteinase K was added, vortexed the suspension, and incubated at room temperature (RT) for 10 min. 2ml of GITC solution was added, and the sample was sonicated for 10 cycles (5 sec/pulse) at 4°C and subjected to continuous vortexing for 15 min. 5ml of Trizol solution was added and incubated for 15 min at RT followed by addition of 1ml chloroform. The mixture was shaken vigorously and incubated on ice for 10 min. The sample was centrifuged, and the aqueous phase was precipitated with 0.5 volume of NH_4_OAc and 2.5 volume of ethanol. Total RNA was quantified by NanoDrop® ND-1000 UV-Vis Spectrophotometer, and quality was checked by electrophoresis on agarose gel. DNA contamination (if any) was removed by DNase I followed by phenol/chloroform extraction.

Total RNA (1µg, from biological replicates samples) was processed individually for cDNA synthesis by AMV reverse transcriptase. Gene-specific primers (2µM) were used for *ppgk* mRNA and 16srRNA element according to standard protocol. Resultant cDNA was purified by QIAquick PCR Purification Kit, concentration was measured by NanoDrop® ND-1000 UV-Vis Spectrophotometer, and ∼10 ng of cDNA was used for quantitative real-time PCR reactions. Quantitative real-time PCR reactions were performed in triplicates by using PowerUp SYBR Green Master Mix (applied biosystems). Reactions were performed in triplicates in 20µl of final reaction volume. Primer concentrations were optimized, and 300nM of primer concentration was used in each reaction. Average Cq value was considered to determine the fold change values.

## Abbreviations

*ppgk*: polyphosphate glucokinase;
SD sequence: Shine Dalgarno sequence

## DECLARATIONS

### Competing interest

Authors declare that they have no competing interest.

### Funding

This work was supported by Council of Scientific and Industrial Research (CSIR), New Delhi, India (Grant no. MLP 6003). Funding body has no role in designing the study, analysis or manuscript preparation.

### Authors Contribution

N.P.B. and I.A.K. conceived the project, and designed the study. N.P.B. performed the experiments, N.P.B. and I.A.K. analysed the results and wrote the manuscript.

## Acknowledgments

We are generously grateful to Prof. Michael Niederweis (University of Alabama, Tuscaloosa, Al, USA) for providing the plasmid pML163 for cloning and expression based assays. Author (Naveen Prakash Bokolia) is thankful to University Grant Commission (UGC), New Delhi, India for providing a Senior Research Fellowship (GAP-1128).

We are very thankful to Dr. Nitin Pal Kalia for helping in the improvement of manuscript and preparation of graphs.

## Table Legend

**Table 1** Determination of 3’RNA fragment by illumina sequencing. Overrepresented counts of the 282-*ppgk* RNA samples transcribed in the presence of 4mM and 8mM Mg^2+^ ions respectively. These counts delivered by overrepresented pattern files, those were extracted by in house python script. R1 and R2 reads are corresponding to reads from both ends of the DNA strand and refers to the top strand (by forward primer) and bottom strand (reverse primer) reads respectively. Overrepresented percent counts were calculated after combining the two respective values, and fractional percentage was calculated. Respective sequences are highlighted in Supplementary data file 1 and 2.

## References

Boniecki MJ, Lach G, Dawson WK, Tomala K, Lukasz P, Soltysinski T, Rother KM, Bujnicki JM. 2015. SimRNA: a coarse-grained method for RNA folding simulations and 3D structure prediction. Nucleic acids research 44: e63–e63.

Brehm SL, Cech TR. 1983. Fate of an intervening sequence ribonucleic acid: excision and cyclization of the Tetrahymena ribosomal ribonucleic acid intervening sequence in vivo. Biochemistry 22: 2390–2397.

Buzayan JM, Gerlach WL, Bruening G. 1986. Non-enzymatic cleavage and ligation of RNAs complementary to a plant virus satellite RNA. Nature 323: 349–353.

Cech TR, Zaug AJ, Grabowski PJ. 1981. In vitro splicing of the ribosomal RNA precursor of Tetrahymena: involvement of a guanosine nucleotide in the excision of the intervening sequence. Cell 27: 487–496.

Chevallet M, Luche S, Rabilloud T. 2006. Silver staining of proteins in polyacrylamide gels. Nat Protoc 1: 1852–1858.

Cook GM, Berney M, Gebhard S, Heinemann M, Cox RA, Danilchanka O, Niederweis M. 2009. Physiology of mycobacteria. Advances in microbial physiology 55: 81–182, 318-189.

Edgar RC. 2004. MUSCLE: multiple sequence alignment with high accuracy and high throughput. Nucleic acids research 32: 1792–1797.

Gabizon R, Lee A, Vahedian-Movahed H, Ebright RH, Bustamante CJ. 2018. Pause sequences facilitate entry into long-lived paused states by reducing RNA polymerase transcription rates. Nature communications 9: 2930–2930.

Hanna R, Doudna JA. 2000. Metal ions in ribozyme folding and catalysis. Current opinion in chemical biology 4: 166–170.

Heilman-Miller SL, Woodson SA. 2003. Effect of transcription on folding of the Tetrahymena ribozyme. RNA (New York, NY) 9: 722–733.

Hofacker IL. 2003. Vienna RNA secondary structure server. Nucleic acids research 31: 3429–3431.

Hsieh PC, Shenoy BC, Samols D, Phillips NF. 1996. Cloning, expression, and characterization of polyphosphate glucokinase from Mycobacterium tuberculosis. The Journal of biological chemistry 271: 4909–4915.

Hubin EA, Fay A, Xu C, Bean JM, Saecker RM, Glickman MS, Darst SA, Campbell EA. 2017. Structure and function of the mycobacterial transcription initiation complex with the essential regulator RbpA. eLife 6.

Incarnato D, Morandi E, Anselmi F, Simon LM, Basile G, Oliviero S. 2017. In vivo probing of nascent RNA structures reveals principles of cotranscriptional folding. Nucleic acids research 45: 9716–9725.

Lai D, Proctor JR, Meyer IM. 2013. On the importance of cotranscriptional RNA structure formation. RNA (New York, NY) 19: 1461–1473.

Li PT, Vieregg J, Tinoco I, Jr. 2008. How RNA unfolds and refolds. Annu Rev Biochem 77: 77–100.

Liu SR, Hu CG, Zhang JZ. 2016. Regulatory effects of cotranscriptional RNA structure formation and transitions. Wiley interdisciplinary reviews RNA 7: 562–574.

Malke H. 1993. Jeffrey H. Miller, A Short Course in Bacterial Genetics–A Laboratory Manual and Handbook for Escherichia coli and Related Bacteria. Cold Spring Harbor 1992. Cold Spring Harbor Laboratory Press. ISBN: 0–87969-349–5. Journal of Basic Microbiology 33: 278–278.

Mandal M, Breaker RR. 2004. Gene regulation by riboswitches. Nature reviews Molecular cell biology 5: 451–463.

Manganelli R, Proveddi R, Rodrigue S, Beaucher J, Gaudreau L, Smith I. 2004. σ Factors and Global Gene Regulation in <em>Mycobacterium tuberculosis</em>. Journal of Bacteriology 186: 895–902.

Nahvi A, Sudarsan N, Ebert MS, Zou X, Brown KL, Breaker RR. 2002. Genetic control by a metabolite binding mRNA. Chemistry & biology 9: 1043.

Nudler E. 2009. RNA polymerase active center: the molecular engine of transcription. Annual review of biochemistry 78: 335–361.

Pan T, Sosnick T. 2006. RNA folding during transcription. Annual review of biophysics and biomolecular structure 35: 161–175.

Perdrizet GA, 2nd, Artsimovitch I, Furman R, Sosnick TR, Pan T. 2012. Transcriptional pausing coordinates folding of the aptamer domain and the expression platform of a riboswitch. Proceedings of the National Academy of Sciences of the United States of America 109: 3323–3328.

Proctor JR, Meyer IM. 2013. COFOLD: an RNA secondary structure prediction method that takes co-transcriptional folding into account. Nucleic acids research 41: e102–e102.

Raman S, Hazra R, Dascher CC, Husson RN. 2004. Transcription Regulation by the <em>Mycobacterium tuberculosis</em> Alternative Sigma Factor SigD and Its Role in Virulence. Journal of Bacteriology 186: 6605–6616.

Regulski EE, Breaker RR. 2008. In-line probing analysis of riboswitches. Methods in molecular biology (Clifton, NJ) 419: 53–67.

Rodriguez GM, Voskuil MI, Gold B, Schoolnik GK, Smith I. 2002. ideR, An essential gene in mycobacterium tuberculosis: role of IdeR in iron-dependent gene expression, iron metabolism, and oxidative stress response. Infection and immunity 70: 3371–3381.

Serganov A, Patel DJ. 2007a. Ribozymes, riboswitches and beyond: regulation of gene expression without proteins. Nature reviews Genetics 8: 776–790.

Serganov A, Patel DJ. 2007b. Ribozymes, riboswitches and beyond: regulation of gene expression without proteins. Nature reviews Genetics 8: 776–790.

Smith I. 2003. Mycobacterium tuberculosis pathogenesis and molecular determinants of virulence. Clin Microbiol Rev 16: 463–496.

Teixeira A, Tahiri-Alaoui A, West S, Thomas B, Ramadass A, Martianov I, Dye M, James W, Proudfoot NJ, Akoulitchev A. 2004. Autocatalytic RNA cleavage in the human beta-globin pre-mRNA promotes transcription termination. Nature 432: 526–530.

Turkarslan S, Peterson EJR, Rustad TR, Minch KJ, Reiss DJ, Morrison R, Ma S, Price ND, Sherman DR, Baliga NS. 2015. A comprehensive map of genome-wide gene regulation in Mycobacterium tuberculosis. Scientific Data 2: 150010.

Wickiser JK, Winkler WC, Breaker RR, Crothers DM. 2005. The speed of RNA transcription and metabolite binding kinetics operate an FMN riboswitch. Molecular cell 18: 49–60.

Winkler W, Nahvi A, Breaker RR. 2002. Thiamine derivatives bind messenger RNAs directly to regulate bacterial gene expression. Nature 419: 952–956.

Winkler WC, Breaker RR. 2005. Regulation of bacterial gene expression by riboswitches. Annual review of microbiology 59: 487–517.

Winkler WC, Nahvi A, Roth A, Collins JA, Breaker RR. 2004. Control of gene expression by a natural metabolite-responsive ribozyme. Nature 428: 281–286.

Wong T, Sosnick TR, Pan T. 2005. Mechanistic insights on the folding of a large ribozyme during transcription. Biochemistry 44: 7535–7542.

Wong TN, Sosnick TR, Pan T. 2007. Folding of noncoding RNAs during transcription facilitated by pausing-induced nonnative structures. Proceedings of the National Academy of Sciences of the United States of America 104: 17995–18000.

Xayaphoummine A, Bucher T, Isambert H. 2005. Kinefold web server for RNA/DNA folding path and structure prediction including pseudoknots and knots. Nucleic acids research 33: W605–610.

Zemora G, Waldsich C. 2010. RNA folding in living cells. RNA biology 7: 634–641.

